## Supplementary material for "Testing Times: Challenges in Disentangling Admixture Histories in Recent and Complex Demographies": Supplementary Material_ Testing Times_ Challenges in Disentangling Admixture Histories in Recent and Complex Demographies.pdf

#### SI Figure S1

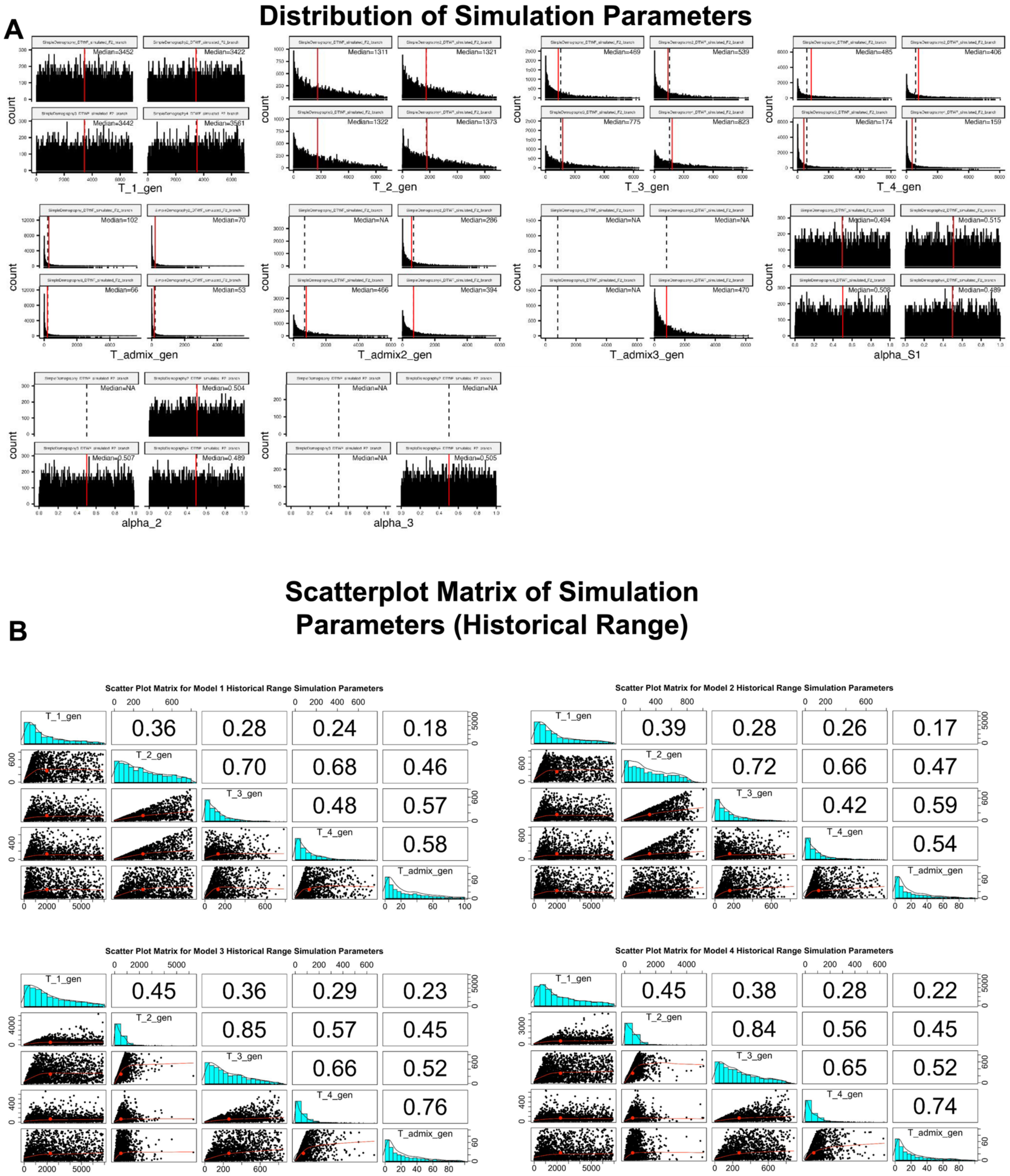


SI Figure 1: The distributions of simulation parameters for the simple demographic Models shown in the Main publication Figure 3A-D. For each simulation parameter the Models are ordered; top left (Model 1), top right

(Model 2), bottom left (Model 3), and bottom right (Model 4). The median value for each simulation parameter within each demographic Model is printed in the top right corner of each box. The median value across all demographic Models for each simulation parameter is represented with the red dotted line. (A) The unconstrained distributions for each of the simulation parameters (T1, T2, T3, T4, Tadmix, Tadmix2, and Tadmix3, alpha_S1, alpha_2, alpha_3). Note that in Model 2, Tadmix2 is the admixture event from the outgroup R3 branch to the S1 population and alpha_2 is its simulated weight parameter. In Model 3, Tadmix2 is the admixture event from the S2R2 branch to S1 and alpha_2 is its simulated weight parameter. The parameter Tadmix 3 is the additional admixture parameter in Model 4 and corresponds to admixture from the S2R2 branch to S1 and alpha_3 is its simulated weight parameter. (B) Scatterplot matrix of simulation parameters (T1, T2, T3, T4, Tadmix) for each demographic Model under the constraint of median pairwise $F_{ST}$ (S1, S2, R1, R2, and R3 populations) between 0 and 0.02 and Tadmix less than or equal to 100 generations. The plots were generated with R package psych using the Spearman correlation method.

#### SI Figure S2

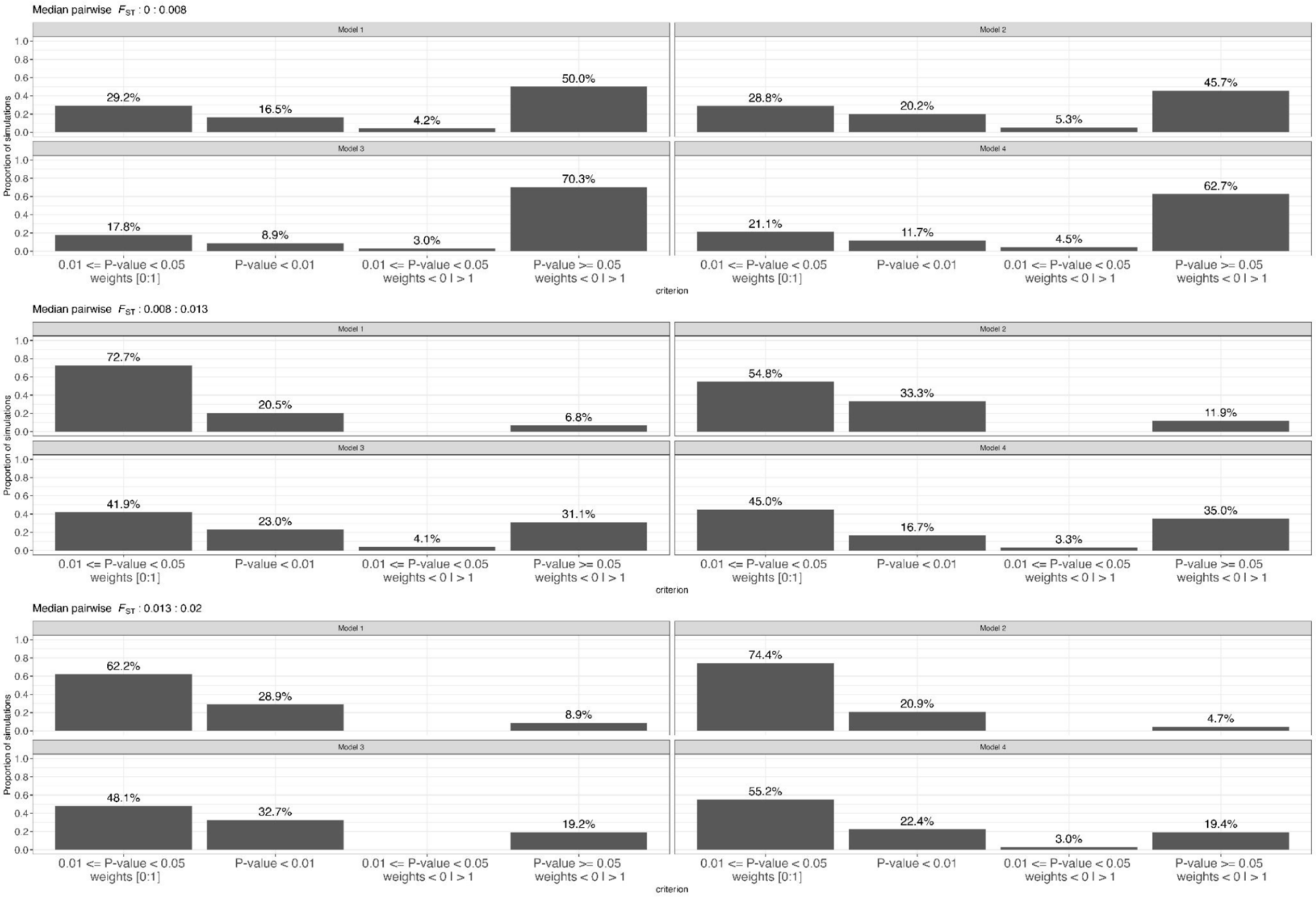


SI Figure 2: The frequency of conditions leading to the rejection of the true qpAdm model (S1+S2) under varying $F_{ST}$ ranges. The Y-axis shows the frequency of that condition across the replicates and the number above each barplot shows the proportion of the total conditions that each makes up.

### SI Figure S3

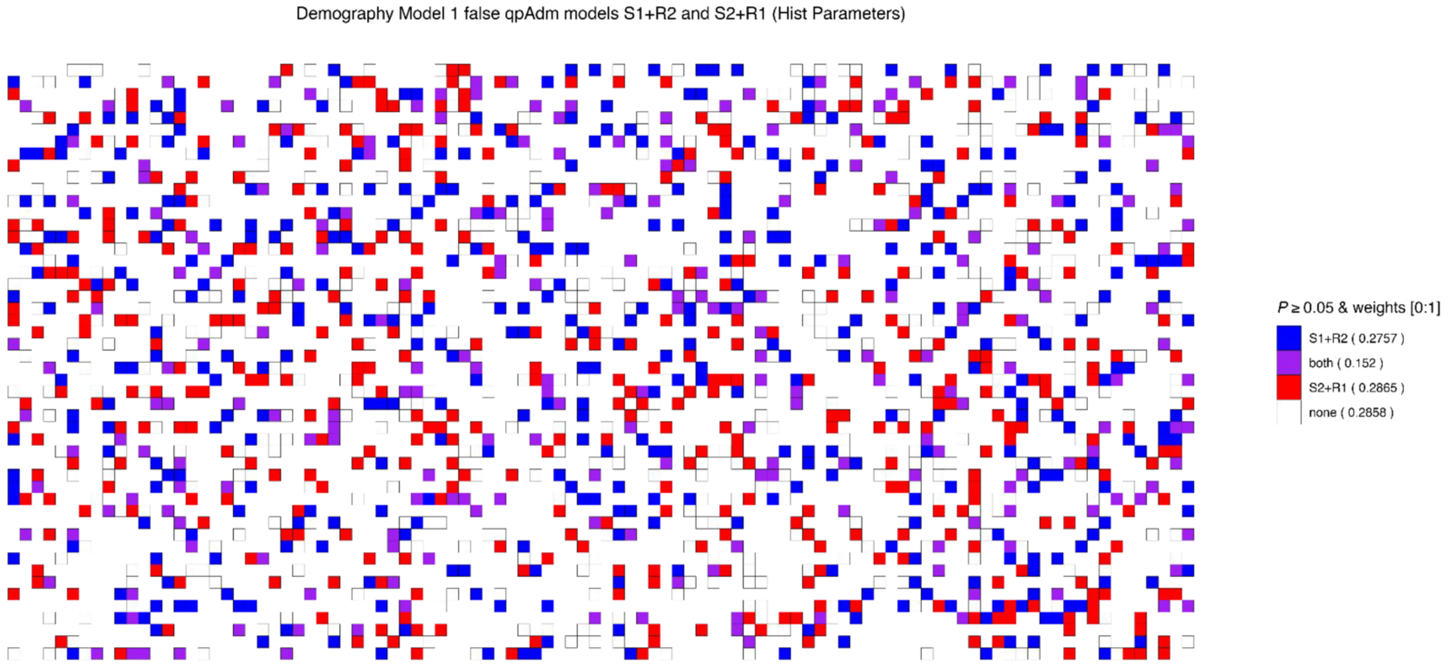


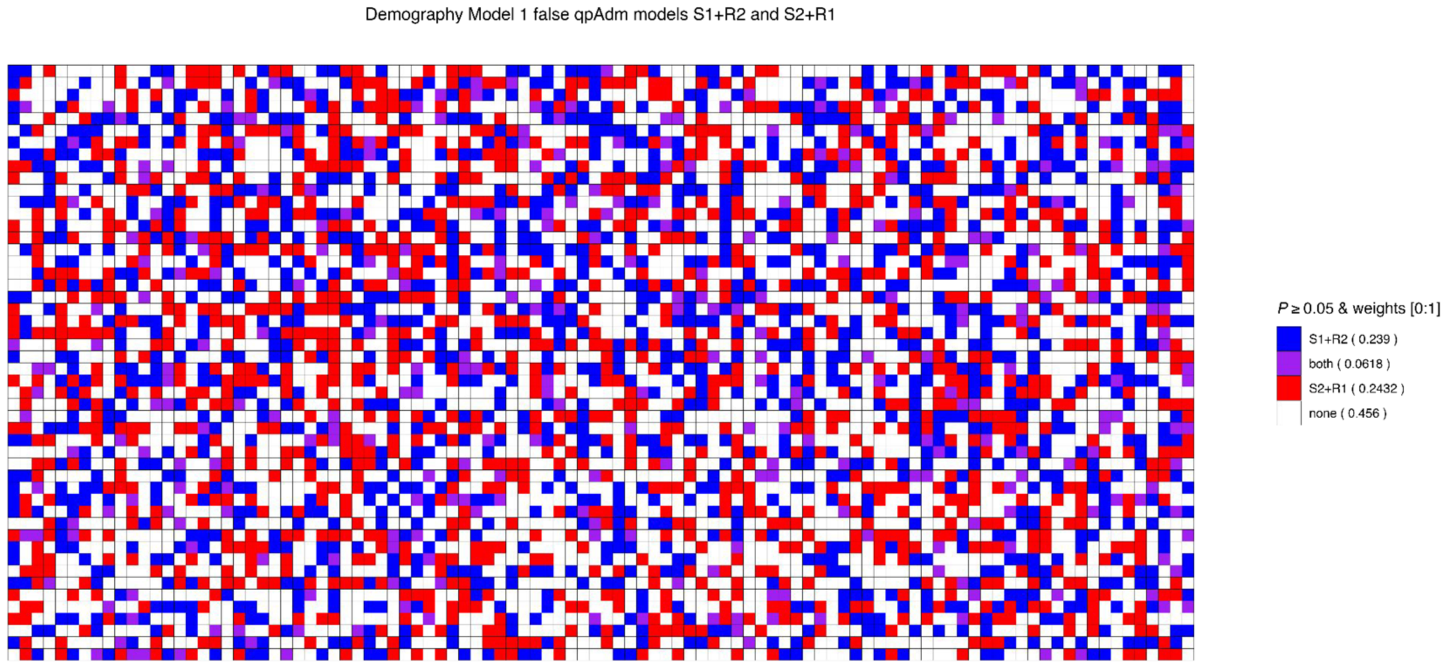


SI Figure 3: Dot matrix plot showing for the historical conditions (top) and full parameter range (bottom) for the cases where in each replicate both false models (S1+R2 & S2+R1) are plausible (purple), or only one of the false models is plausible (red or blue), or both false models are rejected (white).

### SI Figure S4

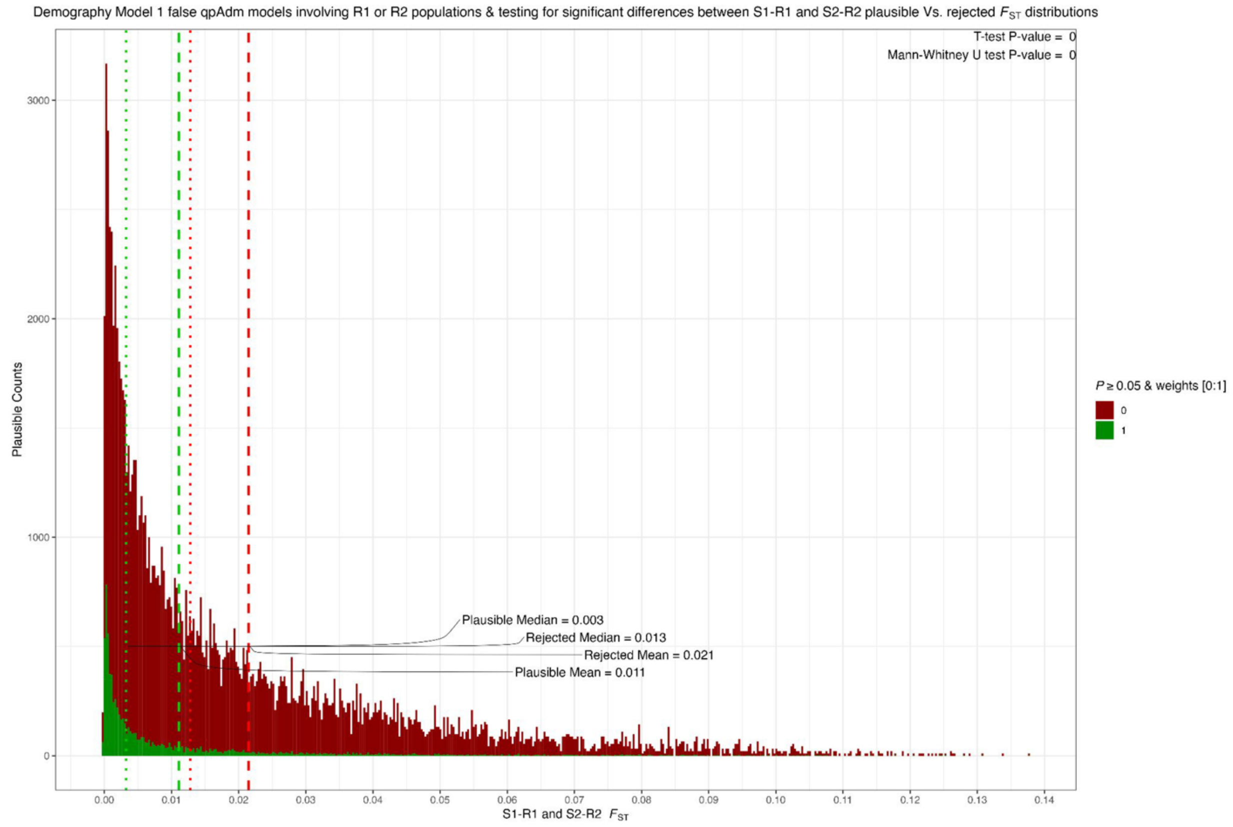


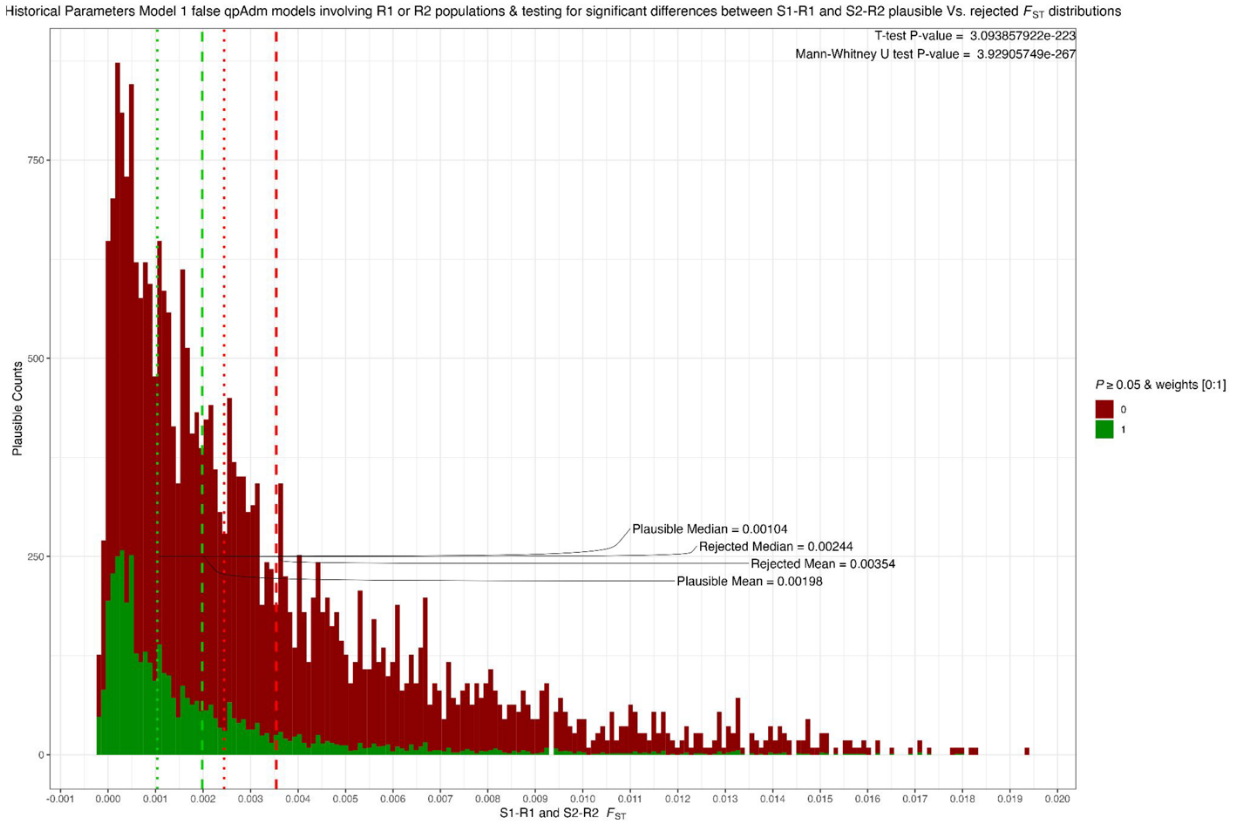


SI Figure 4: The joint distribution of $F_{ST}$ (S1-R1 and S2-R2) for the false qpAdm models involving one of the R1 or R2 populations (Model 1). False models that are plausible have smaller $F_{ST}$ between the clades of S1-R1 and

S2-R2.

SI Figure S5

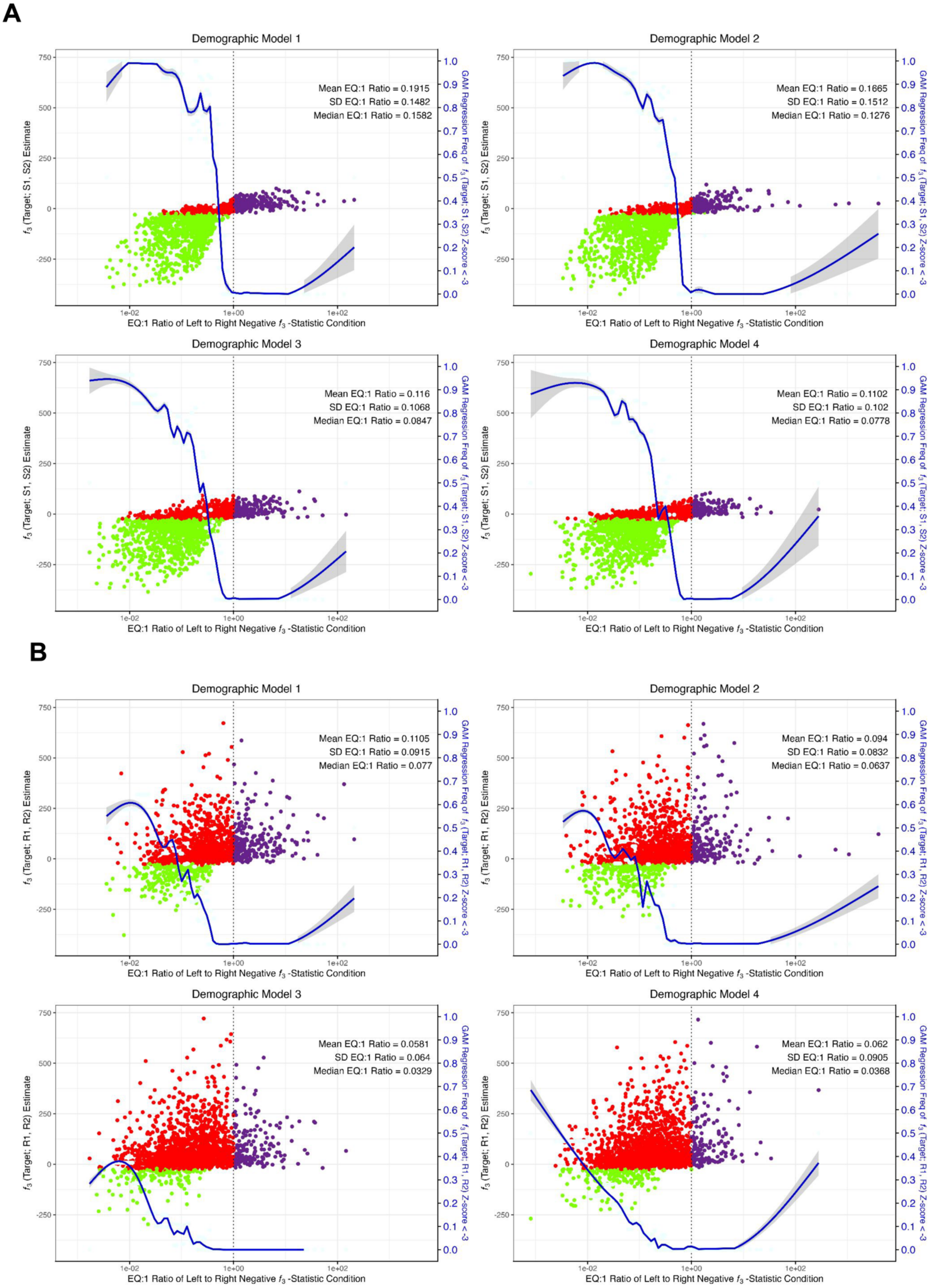

C

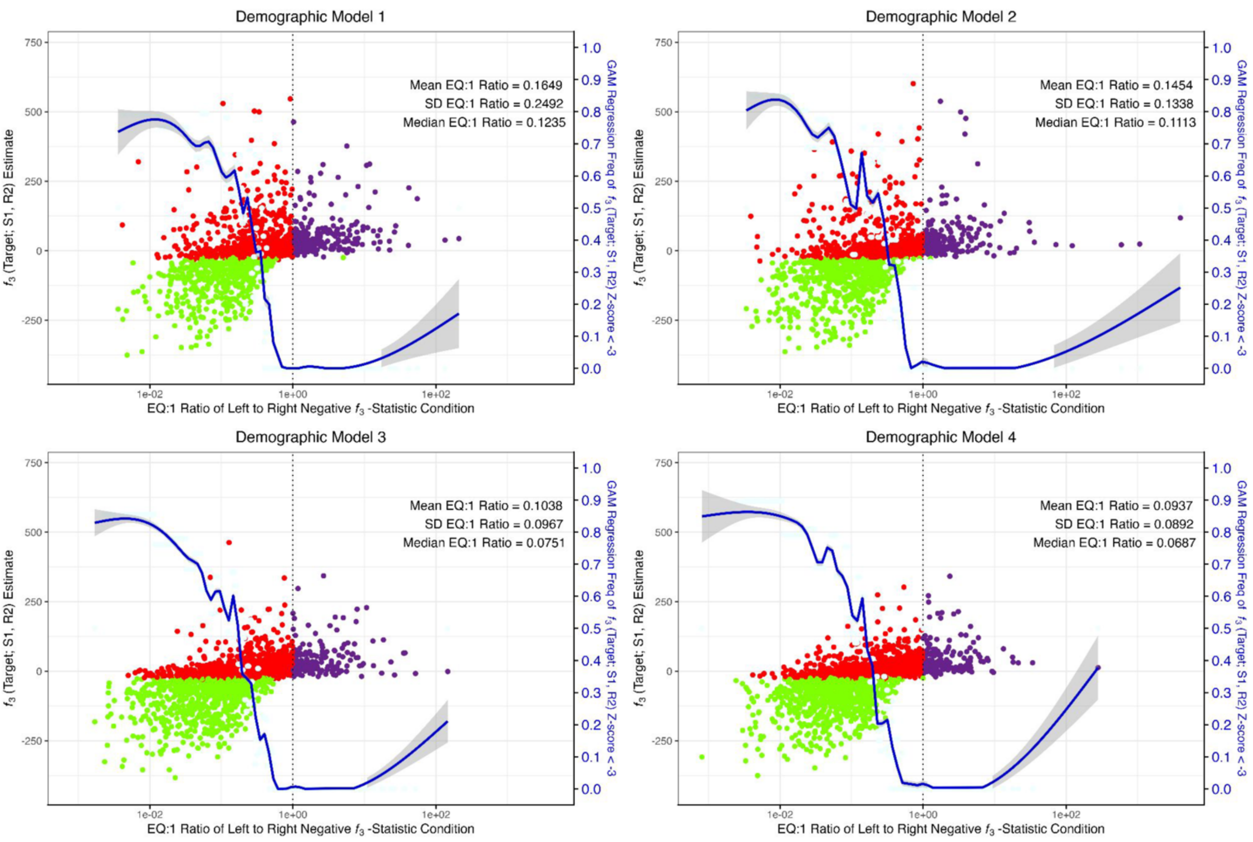


D

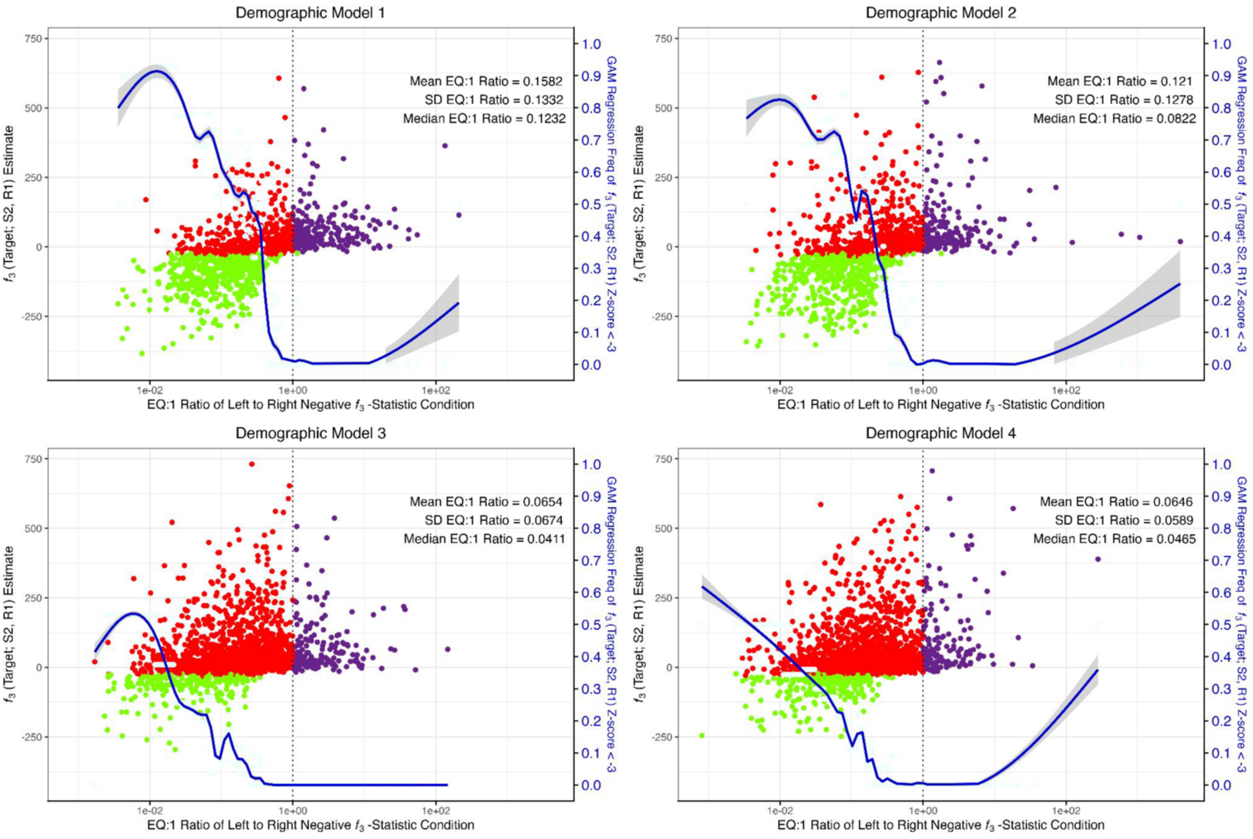


SI Figure 5: The $f_3$-statistic condition, EQ:1 from the manuscript where we are dividing the left side of the condition by the right side (2 alpha (1 - alpha)). Each plot shows on the x axis the EQ:1 $f_3$-statistic condition and on the y axis the $f_3$-statistic estimate. Green points show the $f_3$-statistic estimate with a Z-score < -3, purple points show the $f_3$-statistic estimate with a Z-score > -3, and red points are those with an EQ:1 $f_3$-statistic negativity condition < 1, but the $f_3$-statistic estimate has a Z-score > -3 and thus not significant. In each top right corner of each plot is shown the mean, standard deviation, and median of the EQ:1 negative $f_3$-statistic condition for significant $f_3$-statistics (Z-scores <-3). (A) the $f_3$(Target; S1, S2). (B) $f_3$(Target; R1, R2), (C) $f_3$(Target; S1, R2). (D) $f_3$(Target; S2, R1)

#### SI Figure S6

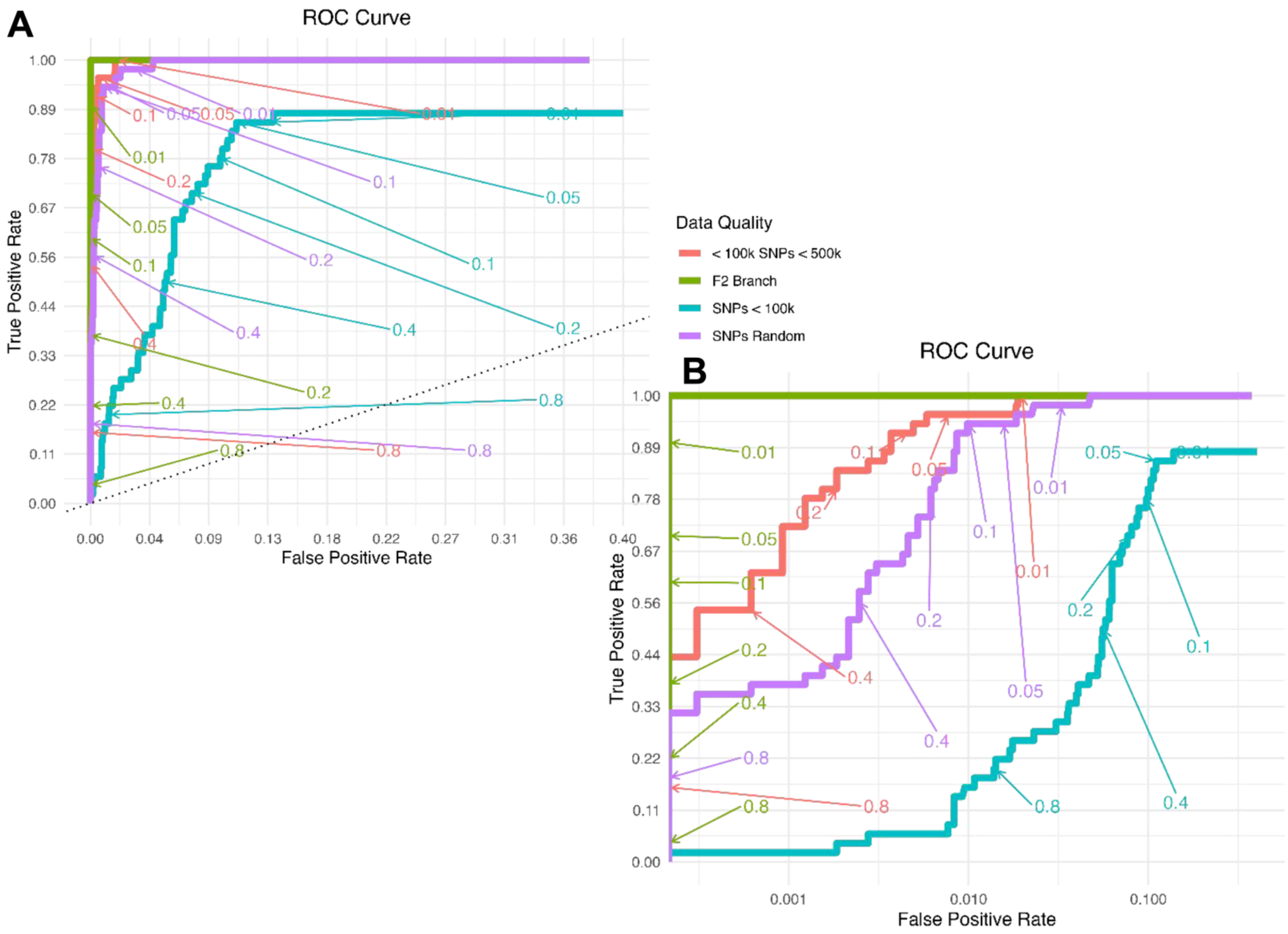


SI Figure 6: Receiver operating characteristic curve (ROC) analysis on *P*-values. Each ROC analysis was computed on 3,000 *P*-values between zero and one. The True Positive Rate is the number of plausible (*P*-value greater than or equal to the iterator value) true qpAdm model observations (TP) divided by the sum of the number of TP and the number of times the true model is rejected (FN) (*P*-value less than the iterator value). The False Positive Rate was calculated similarly, with the number of FP models divided by the sum of FP and true rejections (TN) of single and two-source false qpAdm models. Colors represent different levels of missing data conditions. (A) the ROC analysis across the entire FP range. (B) ROC analysis with the FP range on a log10 scale.

SI Figure S7

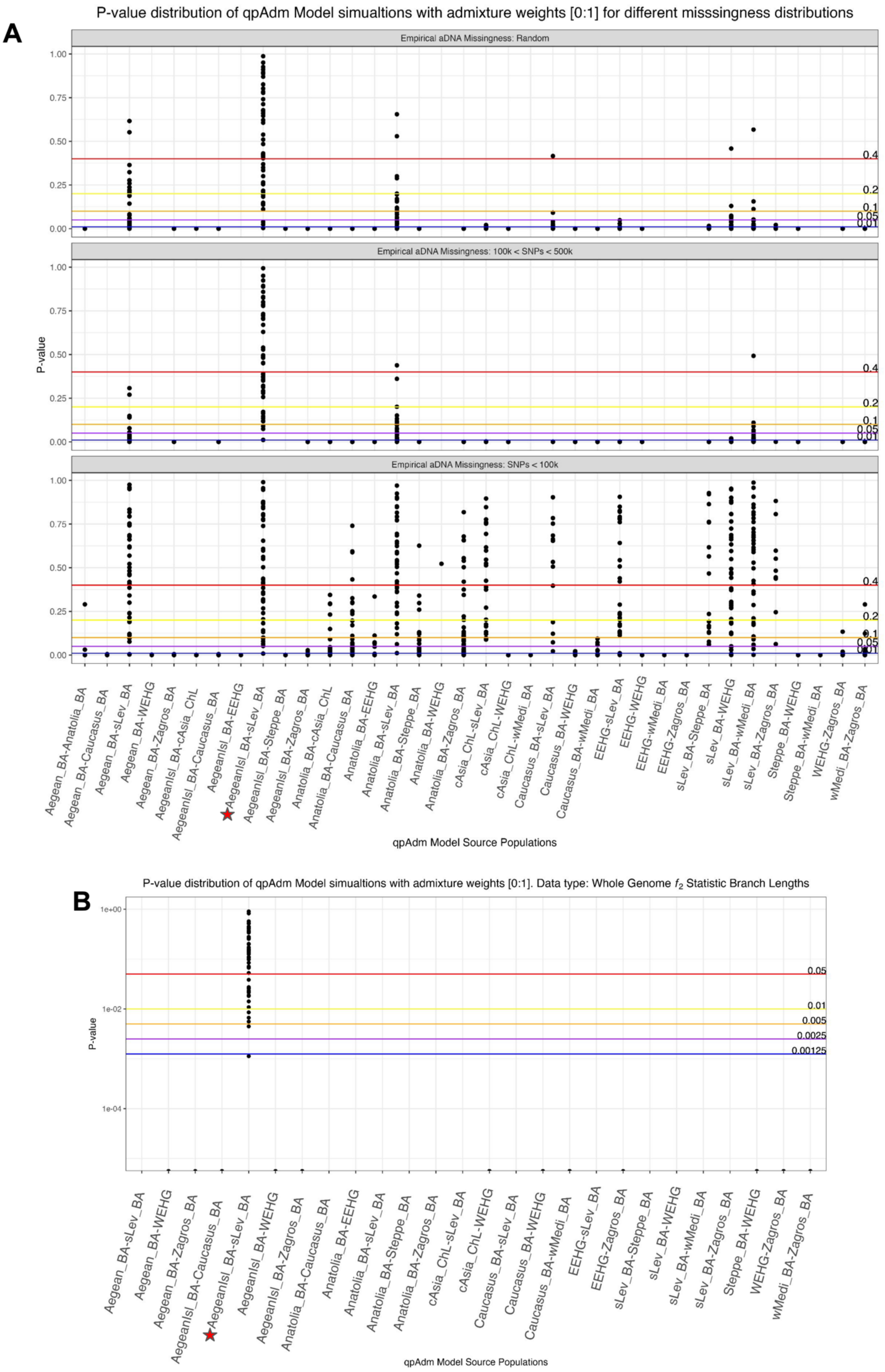


SI Figure 7: Distribution of *P*-values for qpAdm models with admixture weights [0:1]. (A) Distribution of *P*-values for different amounts of data missingness. Horizontal lines show various *P*-values thresholds in the bottom to top order: blue = 0.01, purple = 0.05, orange = 0.1, yellow = 0.2, and red = 0.4. (B) Distribution of *P*-values for the $f_2$ branch data condition. Horizontal lines show various *P*-value thresholds bottom to top order: blue = 0.00125, purple = 0.0025, orange = 0.005, yellow = 0.01, and red = 0.05. Red star below the x axis qpAdm models shows the true model.

#### SI Figure S8

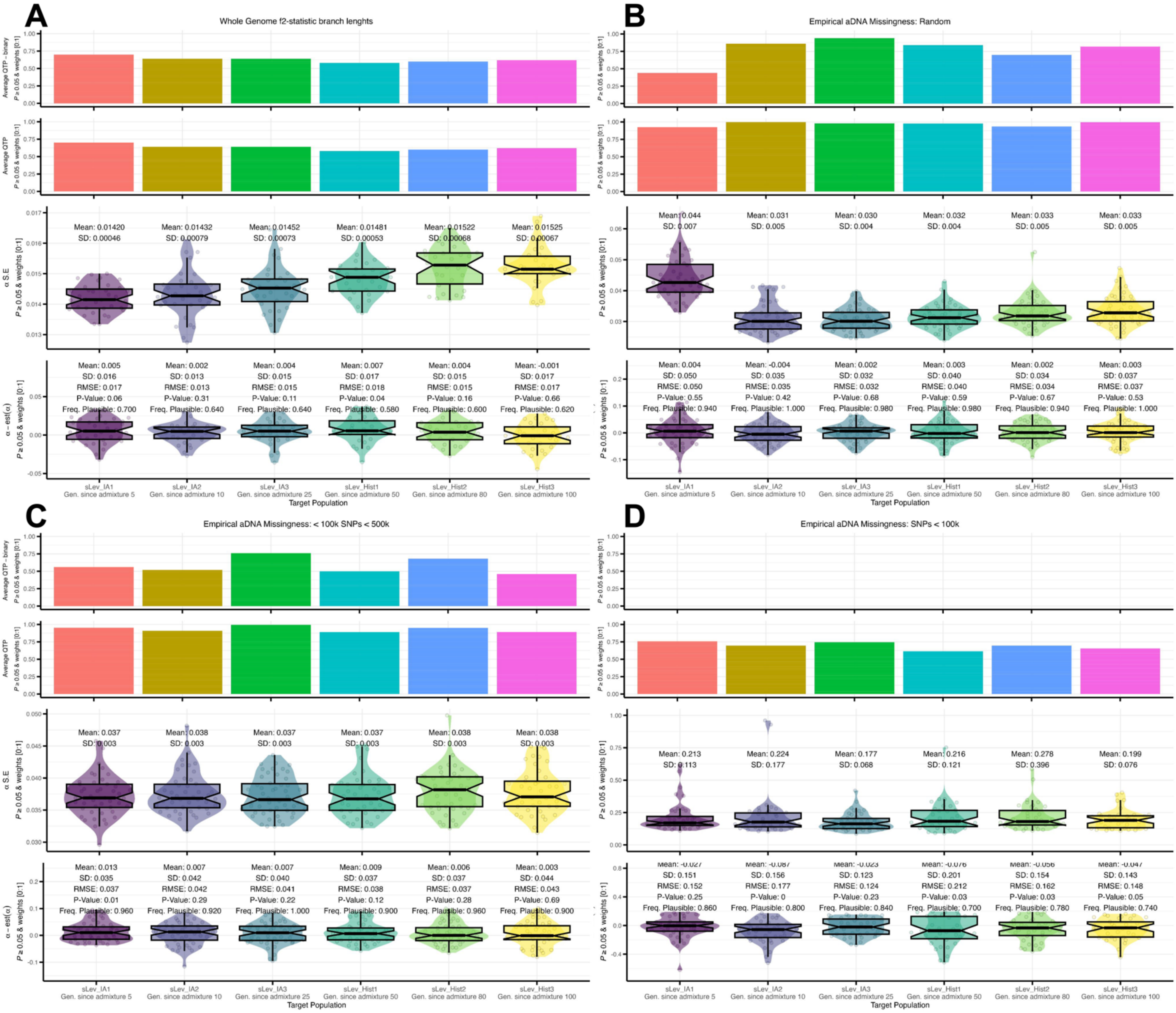


SI Figure 8: Generations since admixture and qpAdm performance (QTP). Measures of qpAdm performance (QTP-binary (top row) and QTP (second row)) averaged across all 50 simulations. The accuracy and precision of qpAdm admixture weight estimates is measured through the average and SD of the estimated standard error across all simulations (middle row) and the average, SD, root-mean-standard error, one-sample t-test *P*-value of the delta-alpha calculation (bottom row). The frequency across all replicates of the plausible true qpAdm model is written at the bottom. (A) Results computed on the branch-length $f_2$-statistic data. (B) Computed on the random sampling of empirical aDNA data missingness (C) Computed on the random sampling of empirical aDNA data missingness constrained to have samples with 100k to 500k SNPs covered. (D) Computed on the random sampling of empirical aDNA data missingness constrained to have samples with less than 100k SNPs covered.

### SI Figure S9

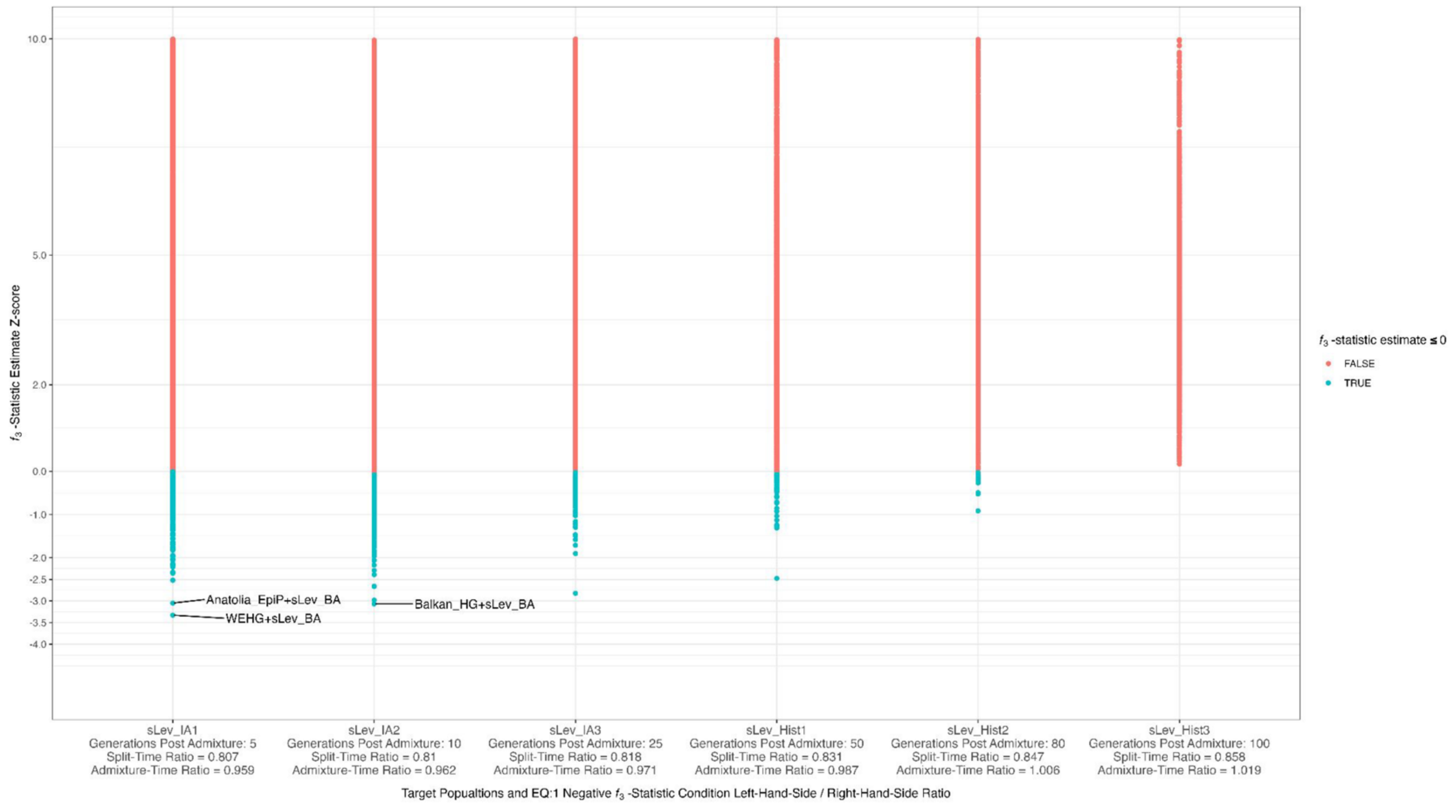


SI Figure S9: The Z-score for the *f3*-statistic estimate computed for the six Target populations each with increasing generations since admixture. $f_3$-statistic values less than zero are green. The two values printed below each Target population are the $f_3$-statistic negativity condition, EQ:1 from the manuscript where we are dividing the left side of the condition by the right side (2 alpha (1 - alpha)). The Tadmix for the Split-time Ratio is computed from the split of the Levant and Aegean populations, and the Admixture-Time Ratio provides the Tadmix with the generation of the most recent bi-directional gene-flow between the Levant and Aegean lineages.

#### SI Figure S10

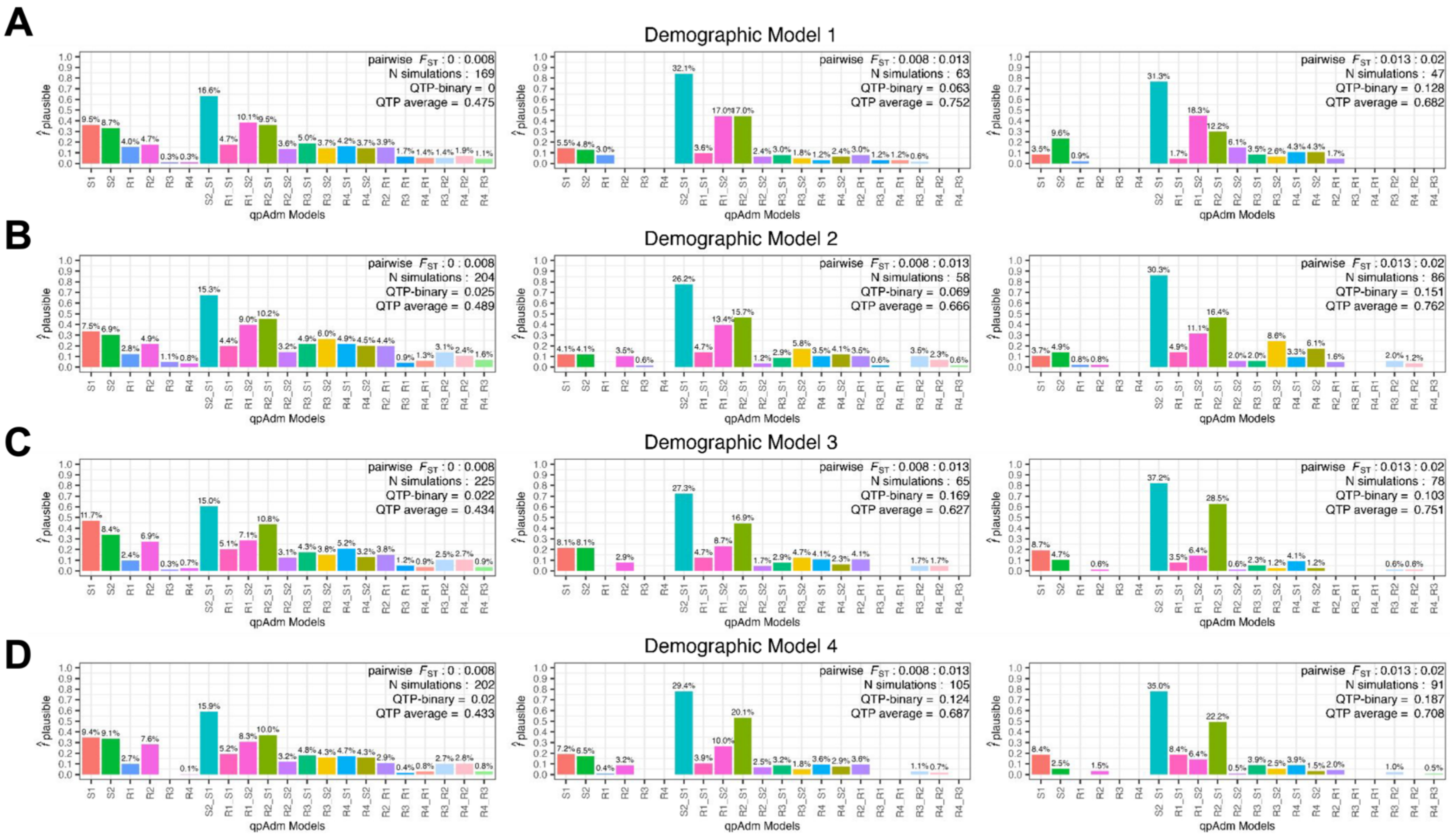


SI Figure S10. Increase in plausible single-source qpAdm models with admixture weights near the boundaries. Barplots of plausible *qpAdm* models conditional on parameter space of the simulated admixture weight < 0.1 or > 0.9, median pairwise $F_{ST}$ between the S1, S2, R1, R2, and R3 populations between 0 and 0.02, and the number of generations since admixture less than 100. Each row represents one simulated demographic history and the columns are increasing ranges of population differentiation ($F_{ST}$) corresponding to the historical period demarcations indicated in Figure 1 B.

#### SI Table ST1

**A**

P-value ranking: Demographic Model 1[1]

| Model_Sources | Freq_Largest_Pvalue | Freq_Second_Largest_Pvalue | Freq_Third_Largest_Pvalue | Freq_Fourth_Largest_Pvalue |
|---|---|---|---|---|
| S2_S1 | 0.654 | 0.221 | 0.019 | 0.014 |
| R1_S2 | 0.137 | 0.151 | 0.024 | 0.012 |
| R2_S1 | 0.127 | 0.148 | 0.029 | 0.012 |
| R2_R1 | 0.011 | 0.014 | 0.017 | 0.022 |
| R1_S1 | 0.011 | 0.004 | 0.004 | 0.002 |
| S2 | 0.008 | 0.008 | 0.012 | 0.005 |
| R2_S2 | 0.007 | 0.005 | 0.005 | 0.004 |
| S1 | 0.007 | 0.010 | 0.012 | 0.006 |
| R3_S1 | 0.006 | 0.004 | 0.003 | 0.006 |
| R4_R3 | 0.006 | 0.003 | 0.001 | 0.000 |
| R1 | 0.005 | 0.005 | 0.005 | 0.004 |
| R2 | 0.004 | 0.004 | 0.005 | 0.004 |
| R4_S2 | 0.003 | 0.003 | 0.003 | 0.004 |
| R4_S1 | 0.003 | 0.003 | 0.005 | 0.002 |
| R3_R2 | 0.002 | 0.001 | 0.001 | 0.002 |
| R3_S2 | 0.002 | 0.003 | 0.005 | 0.004 |
| R3_R1 | 0.002 | 0.002 | 0.002 | 0.002 |
| R4_R1 | 0.002 | 0.001 | 0.001 | 0.002 |
| R4_R2 | 0.001 | 0.001 | 0.001 | 0.001 |
| R3 | 0.000 | 0.001 | 0.000 | 0.000 |
| R4 | 0.000 | 0.000 | 0.001 | 0.000 |

[1] Table of plausible (P-value 0.05 + weights [0:1]) qpAdm models showing the frequency with which each qpAdm model's P-value ranked from the largest to the fourth largest

**B**

P-value ranking: Demographic Model 2[1]

| Model_Sources | Freq_Largest_Pvalue | Freq_Second_Largest_Pvalue | Freq_Third_Largest_Pvalue | Freq_Fourth_Largest_Pvalue |
|---|---|---|---|---|
| S2_S1 | 0.688 | 0.184 | 0.024 | 0.014 |
| R2_S1 | 0.154 | 0.166 | 0.022 | 0.012 |
| R1_S2 | 0.070 | 0.086 | 0.024 | 0.013 |
| R1_S1 | 0.011 | 0.003 | 0.007 | 0.006 |
| R2_R1 | 0.010 | 0.010 | 0.014 | 0.015 |
| R3_S2 | 0.010 | 0.015 | 0.013 | 0.007 |
| R4_S1 | 0.010 | 0.005 | 0.008 | 0.004 |
| R4_S2 | 0.009 | 0.004 | 0.004 | 0.008 |
| S2 | 0.006 | 0.010 | 0.010 | 0.005 |
| R2_S2 | 0.005 | 0.004 | 0.005 | 0.002 |
| S1 | 0.005 | 0.007 | 0.011 | 0.005 |
| R3_S1 | 0.004 | 0.004 | 0.004 | 0.003 |
| R2 | 0.003 | 0.005 | 0.004 | 0.005 |
| R1 | 0.003 | 0.005 | 0.004 | 0.002 |
| R3_R2 | 0.003 | 0.002 | 0.004 | 0.002 |
| R4_R3 | 0.003 | 0.002 | 0.003 | 0.002 |
| R4_R2 | 0.002 | 0.003 | 0.002 | 0.003 |
| R4_R1 | 0.002 | 0.001 | 0.002 | 0.002 |
| R3_R1 | 0.001 | 0.001 | 0.000 | 0.001 |
| R4 | 0.001 | 0.001 | 0.001 | 0.001 |
| R3 | 0.000 | 0.001 | 0.001 | 0.001 |

[1] Table of plausible (P-value 0.05 + weights [0:1]) qpAdm models showing the frequency with which each qpAdm model's P-value ranked from the largest to the fourth largest

**C**

P-value ranking: Demographic Model 3[1]

| Model_Sources | Freq_Largest_Pvalue | Freq_Second_Largest_Pvalue | Freq_Third_Largest_Pvalue | Freq_Fourth_Largest_Pvalue |
|---|---|---|---|---|
| S2_S1 | 0.624 | 0.225 | 0.026 | 0.012 |
| R2_S1 | 0.240 | 0.257 | 0.028 | 0.009 |
| R1_S2 | 0.024 | 0.030 | 0.019 | 0.011 |
| S1 | 0.021 | 0.023 | 0.025 | 0.013 |
| R1_S1 | 0.021 | 0.013 | 0.015 | 0.007 |
| S2 | 0.012 | 0.012 | 0.010 | 0.009 |
| R4_S1 | 0.010 | 0.007 | 0.006 | 0.009 |
| R2_R1 | 0.009 | 0.009 | 0.010 | 0.011 |
| R2 | 0.008 | 0.009 | 0.008 | 0.009 |
| R3_S1 | 0.007 | 0.007 | 0.007 | 0.007 |
| R2_S2 | 0.007 | 0.005 | 0.005 | 0.002 |
| R4_S2 | 0.003 | 0.003 | 0.002 | 0.005 |
| R3_S2 | 0.003 | 0.003 | 0.004 | 0.005 |
| R1 | 0.003 | 0.006 | 0.008 | 0.004 |
| R4_R3 | 0.003 | 0.002 | 0.002 | 0.002 |
| R3_R1 | 0.001 | 0.000 | 0.001 | 0.001 |
| R3_R2 | 0.001 | 0.002 | 0.003 | 0.002 |
| R4_R1 | 0.001 | 0.002 | 0.001 | 0.001 |
| R4_R2 | 0.001 | 0.002 | 0.002 | 0.002 |
| R4 | 0.000 | 0.000 | 0.000 | 0.000 |
| R3 | 0.000 | 0.000 | 0.000 | 0.000 |

[1] Table of plausible (P-value 0.05 + weights [0:1]) qpAdm models showing the frequency with which each qpAdm model's P-value ranked from the largest to the fourth largest

**D**

P-value ranking: Demographic Model 4[1]

| Model_Sources | Freq_Largest_Pvalue | Freq_Second_Largest_Pvalue | Freq_Third_Largest_Pvalue | Freq_Fourth_Largest_Pvalue |
|---|---|---|---|---|
| S2_S1 | 0.615 | 0.235 | 0.023 | 0.013 |
| R2_S1 | 0.257 | 0.254 | 0.026 | 0.008 |
| S1 | 0.022 | 0.020 | 0.021 | 0.007 |
| R1_S2 | 0.021 | 0.030 | 0.019 | 0.013 |
| R1_S1 | 0.016 | 0.012 | 0.014 | 0.006 |
| S2 | 0.015 | 0.014 | 0.011 | 0.006 |
| R2_R1 | 0.009 | 0.011 | 0.009 | 0.011 |
| R4_S1 | 0.008 | 0.006 | 0.006 | 0.005 |
| R2_S2 | 0.007 | 0.007 | 0.003 | 0.003 |
| R3_S1 | 0.006 | 0.007 | 0.004 | 0.011 |
| R2 | 0.005 | 0.010 | 0.009 | 0.007 |
| R3_S2 | 0.004 | 0.004 | 0.006 | 0.004 |
| R4_S2 | 0.004 | 0.004 | 0.004 | 0.004 |
| R1 | 0.003 | 0.005 | 0.004 | 0.005 |
| R4_R3 | 0.003 | 0.002 | 0.002 | 0.001 |
| R3_R2 | 0.002 | 0.002 | 0.002 | 0.002 |
| R4_R2 | 0.002 | 0.001 | 0.002 | 0.001 |
| R3_R1 | 0.001 | 0.000 | 0.001 | 0.001 |
| R3 | 0.000 | 0.000 | 0.000 | 0.000 |
| R4_R1 | 0.000 | 0.001 | 0.002 | 0.001 |
| R4 | 0.000 | 0.000 | 0.000 | 0.000 |

[1] Table of plausible (P-value 0.05 + weights [0:1]) qpAdm models showing the frequency with which each qpAdm model's P-value ranked from the largest to the fourth largest

SI Table ST1: Relative ranking of plausible qpAdm models by *P*-value for the four simple demographic Models.

#### SI Table ST2

*Simulated Data Missingness*

| pop | mu_missing | max_missi ng | min_missing | sd_missing |
|---|---|---|---|---|

| | | | | |
|---|---|---|---|---|
| sLev_IA1 | 0.89 | 0.95 | 0.76 | 0.05 |
| sLev_IA2 | 0.51 | 0.89 | 0.32 | 0.14 |
| sLev_IA3 | 0.49 | 0.69 | 0.36 | 0.08 |
| sLev_Hist1 | 0.58 | 0.78 | 0.38 | 0.09 |
| sLev_Hist2 | 0.63 | 0.76 | 0.47 | 0.08 |
| sLev_Hist3 | 0.60 | 0.91 | 0.35 | 0.12 |
| ascertain_CEU | 0.00 | 0.00 | 0.00 | 0.00 |
| ascertain_CHB | 0.00 | 0.00 | 0.00 | 0.00 |
| ascertain_AFR | 0.00 | 0.00 | 0.00 | 0.00 |
| ascertain_sAs | 0.00 | 0.00 | 0.00 | 0.00 |
| Mbuti | 0.00 | 0.00 | 0.00 | 0.00 |
| UstIshim | 0.63 | 0.86 | 0.39 | 0.10 |
| nEur_HG | 0.73 | 0.92 | 0.44 | 0.10 |
| nAfr_EpiP | 0.78 | 0.92 | 0.54 | 0.09 |
| sLev_EpiP | 0.76 | 0.91 | 0.52 | 0.09 |
| Zagros_Neo | 0.67 | 0.87 | 0.43 | 0.10 |
| Balkan_HG | 0.72 | 0.92 | 0.53 | 0.09 |
| Caucasus_HG | 0.73 | 0.89 | 0.54 | 0.08 |
| Anatolia_EpiP | 0.73 | 0.93 | 0.56 | 0.08 |

| | | | | |
|---|---|---|---|---|
| sLevant_Neo | 0.78 | 0.99 | 0.55 | 0.12 |
| Caucasus_BA | 0.86 | 0.99 | 0.26 | 0.13 |
| Zagros_BA | 0.86 | 0.99 | 0.36 | 0.15 |
| cAsia_ChL | 0.65 | 0.99 | 0.31 | 0.21 |
| wMedi_BA | 0.45 | 0.81 | 0.27 | 0.12 |
| sLev_BA | 0.51 | 0.81 | 0.28 | 0.14 |
| Aegean_BA | 0.64 | 0.90 | 0.38 | 0.14 |
| AegeanIsl_BA | 0.69 | 0.91 | 0.40 | 0.12 |
| Steppe_BA | 0.64 | 0.92 | 0.44 | 0.13 |
| WEHG | 0.55 | 0.86 | 0.33 | 0.11 |
| EEHG | 0.49 | 0.72 | 0.27 | 0.10 |
| Anatolia_BA | 0.46 | 0.66 | 0.28 | 0.09 |
| BasalEurasian | 0.52 | 0.65 | 0.29 | 0.08 |
| eEurasia | 0.54 | 0.74 | 0.36 | 0.08 |
| nEurasia | 0.54 | 0.86 | 0.34 | 0.13 |
| OOA | 0.77 | 0.95 | 0.33 | 0.19 |
| NearEast | 0.73 | 0.95 | 0.33 | 0.19 |
| eNearEast | 0.68 | 0.96 | 0.08 | 0.19 |
| wNearEast | 0.85 | 0.99 | 0.42 | 0.13 |

| | | | | |
|---|---|---|---|---|
| nwNearEast | 0.53 | 0.77 | 0.35 | 0.10 |

*Note. Missingness from the random sampling scheme of Southwest Asian individuals from Allen Ancient DNA Resource (AADR) v.52.2.*

#### SI Table ST3

| qpAdm Average Performance: Eurasian Simulations & Data Quality | | | | | | | | | | | | | | | | |
|---|---|---|---|---|---|---|---|---|---|---|---|---|---|---|---|---|
| | WG Branch-Length F2-statistics | | | | aDNA Random Sampling | | | | aDNA 100k < SNPs < 500k | | | | aDNA SNPs < 100k | | | |
| **Plausibility Criteria** | FP | FDR | QTP | QTP-binary | FP | FDR | QTP | QTP-binary | FP | FDR | QTP | QTP-binary | FP | FDR | QTP | QTP-binary |
| *P-value 0.01* | 0.000 | 0.000 | 0.900 | 0.900 | 0.044 | 0.530 | 0.936 | 0.240 | 0.027 | 0.464 | 0.973 | 0.280 | 0.344 | 0.958 | 0.616 | 0.000 |
| *P-value 0.05* | 0.000 | 0.000 | 0.700 | 0.700 | 0.021 | 0.345 | 0.919 | 0.420 | 0.009 | 0.239 | 0.951 | 0.560 | 0.262 | 0.945 | 0.698 | 0.000 |
| *P-value 0.01 + weights [0:1]* | 0.000 | 0.000 | 0.900 | 0.900 | 0.033 | 0.511 | 0.947 | 0.240 | 0.020 | 0.428 | 0.980 | 0.300 | 0.139 | 0.907 | 0.721 | 0.000 |
| *P-value 0.05 + weights [0:1]* | 0.000 | 0.000 | 0.700 | 0.700 | 0.016 | 0.328 | 0.924 | 0.440 | 0.008 | 0.226 | 0.952 | 0.560 | 0.112 | 0.881 | 0.748 | 0.000 |
| *P-value 0.01 + weights [0:1] ± 2s.e* | 0.000 | 0.000 | 0.900 | 0.900 | 0.023 | 0.492 | 0.917 | 0.240 | 0.019 | 0.422 | 0.981 | 0.300 | 0.055 | 0.971 | 0.105 | 0.000 |
| *P-value 0.05 + weights [0:1] ± 2s.e* | 0.000 | 0.000 | 0.700 | 0.700 | 0.011 | 0.311 | 0.889 | 0.460 | 0.008 | 0.226 | 0.952 | 0.560 | 0.039 | 0.957 | 0.121 | 0.000 |
| *All single-source models rejected* | | | | | | | | | | | | | | | | |
| *P-value 0.01 + weights [0:1]* | 0.000 | 0.000 | | | 0.019 | 0.539 | | | 0.019 | 0.431 | | | 0.022 | 1.000 | | |
| *P-value 0.05 + weights [0:1]* | 0.000 | 0.000 | | | 0.012 | 0.339 | | | 0.008 | 0.226 | | | 0.022 | 0.965 | | |
| *P-value 0.01 + weights [0:1] ± 2s.e* | 0.000 | 0.000 | | | 0.018 | 0.530 | | | 0.018 | 0.426 | | | 0.022 | 1.000 | | |
| *P-value 0.05 + weights [0:1] ± 2s.e* | 0.000 | 0.000 | | | 0.010 | 0.317 | | | 0.008 | 0.226 | | | 0.020 | 0.981 | | |
| *All single-source models rejected & significant f3-statistics* | | | | | | | | | | | | | | | | |
| *P-value 0.01 + weights [0:1]* | 0.000 | | | | 0.003 | 1.000 | | | 0.000 | 1.000 | | | 0.022 | 1.000 | | |
| *P-value 0.05 + weights [0:1]* | 0.000 | | | | 0.001 | 1.000 | | | 0.000 | | | | 0.015 | 1.000 | | |
| *P-value 0.01 + weights [0:1] ± 2s.e* | 0.000 | | | | 0.003 | 1.000 | | | 0.000 | 1.000 | | | 0.022 | 1.000 | | |
| *P-value 0.05 + weights [0:1] ± 2s.e* | 0.000 | | | | 0.001 | 1.000 | | | 0.000 | | | | 0.015 | 1.000 | | |

SI Table ST3: Performance summaries of rotating qpAdm analysis on the West Eurasian simulations for the four datasets differing in data missingness levels and for different performance metrics: FP = false positive rate, FDR = false discovery rate, QTP = qpAdm test performance, QTP-binary = qpAdm test performance provided that only the true model fits the data.

### SI Table ST4

**A**

P-value ranking: Empirical aDNA Missingness: SNPs < 100k[1]

| Model_Sources | Freq_Largest_Pvalue | Freq_Second_Largest_Pvalue | Freq_Third_Largest_Pvalue | Freq_Fourth_Largest_Pvalue |
|---|---|---|---|---|
| sLev_BA+Anatolia_BA | 0.20 | 0.16 | 0.10 | 0.08 |
| ★ sLev_BA+AegeanIsl_BA | 0.16 | 0.18 | 0.12 | 0.14 |
| wMedi_BA+sLev_BA | 0.14 | 0.10 | 0.12 | 0.08 |
| sLev_BA | 0.10 | 0.08 | 0.12 | 0.14 |
| Caucasus_BA+sLev_BA | 0.06 | 0.00 | 0.02 | 0.02 |
| Zagros_BA+Anatolia_BA | 0.06 | 0.08 | 0.04 | 0.02 |
| Zagros_BA+sLev_BA | 0.06 | 0.00 | 0.04 | 0.02 |
| sLev_BA+Aegean_BA | 0.06 | 0.12 | 0.12 | 0.10 |
| cAsia_ChL+sLev_BA | 0.04 | 0.00 | 0.02 | 0.08 |
| sLev_BA+WEHG | 0.04 | 0.12 | 0.12 | 0.10 |
| Caucasus_BA+Anatolia_BA | 0.02 | 0.06 | 0.08 | 0.02 |
| Steppe_BA+Anatolia_BA | 0.02 | 0.00 | 0.00 | 0.00 |
| sLev_BA+EEHG | 0.02 | 0.04 | 0.02 | 0.04 |
| sLev_BA+Steppe_BA | 0.02 | 0.04 | 0.00 | 0.02 |
| Caucasus_BA+wMedi_BA | 0.00 | 0.00 | 0.00 | 0.02 |
| WEHG+Anatolia_BA | 0.00 | 0.00 | 0.02 | 0.00 |
| Zagros_BA+wMedi_BA | 0.00 | 0.00 | 0.02 | 0.00 |
| cAsia_ChL+Anatolia_BA | 0.00 | 0.00 | 0.00 | 0.02 |

[1] Table of plausible (P-value 0.05 + weights [0:1]) qpAdm models showing the frequency with which each qpAdm model's P-value ranked from the largest to the fourth largest

**B**

P-value ranking: Empirical aDNA Missingness: Random[1]

| Model_Sources | Freq_Largest_Pvalue | Freq_Second_Largest_Pvalue | Freq_Third_Largest_Pvalue | Freq_Fourth_Largest_Pvalue |
|---|---|---|---|---|
| ★ sLev_BA+AegeanIsl_BA | 0.936 | 0.064 | 0.000 | 0.000 |
| sLev_BA+Anatolia_BA | 0.043 | 0.149 | 0.149 | 0.021 |
| sLev_BA+Aegean_BA | 0.021 | 0.277 | 0.043 | 0.043 |
| Caucasus_BA+sLev_BA | 0.000 | 0.000 | 0.021 | 0.000 |
| sLev_BA | 0.000 | 0.000 | 0.021 | 0.000 |
| sLev_BA+WEHG | 0.000 | 0.000 | 0.021 | 0.000 |
| wMedi_BA+sLev_BA | 0.000 | 0.043 | 0.021 | 0.043 |

[1] Table of plausible (P-value 0.05 + weights [0:1]) qpAdm models showing the frequency with which each qpAdm model's P-value ranked from the largest to the fourth largest

**C**

P-value ranking: Empirical aDNA Missingness: 100k < SNPs < 500k[1]

| Model_Sources | Freq_Largest_Pvalue | Freq_Second_Largest_Pvalue | Freq_Third_Largest_Pvalue | Freq_Fourth_Largest_Pvalue |
|---|---|---|---|---|
| ★ sLev_BA+AegeanIsl_BA | 1 | 0.000 | 0.000 | NaN |
| sLev_BA+Aegean_BA | 0 | 0.146 | 0.000 | NaN |
| sLev_BA+Anatolia_BA | 0 | 0.229 | 0.042 | NaN |
| wMedi_BA+sLev_BA | 0 | 0.042 | 0.062 | NaN |

[1] Table of plausible (P-value 0.05 + weights [0:1]) qpAdm models showing the frequency with which each qpAdm model's P-value ranked from the largest to the fourth largest

SI Table ST4: Relative ranking of plausible qpAdm models by *P*-value. A red star is placed next to the true model. (A) Lowest-coverage aDNA dataset. (B) Random sampling of empirical missingness. (C) Constraint to individuals with 100k to 500k SNPs covered.
